# Mathematical modeling using box-behnken response surface design for optimal DNA yield with variable parameters of time, temperature, and proteinase K concentration

**DOI:** 10.1101/2024.07.21.604461

**Authors:** Farwa Afzal, Saeeda Zia, Rashid Saif

## Abstract

The current research aims to explore an enzyme-assisted inorganic DNA extraction protocol to obtain maximum yield from whole blood. We considered various operating conditions such as incubation temperature and time along with proteinase K enzyme concentration. To model and simulate the extraction process, Box-Behnken Design (BBD) was employed with three factors and its five levels, coupled with the desired functional response surface methodology (RSM). RSM entails fitting a mathematical model for experimental data to optimize a response variable with Box-Behnken Design. The best simulation settings yielded the maximum DNA at an incubation temperature of 56°C for 10 hours and proteinase K enzyme concentration of 10*µ*L. Utilizing these parameters, a yield of 300*µ*L was obtained. Furthermore, other DNA extraction parameters may also be added for a more robust design for obtaining the maximum DNA yield.

## 1 Introduction

In contemporary biomedical research and diagnostics, the extraction of DNA from whole blood specimens is a crucial procedure with extensive applications ranging from genetic analysis to disease detection. As the demand for efficiency and reliability in extraction methodologies increases, researchers are exploring innovative approaches utilizing materials such as magnetic nanoparticles and silica matrices to overcome the limitations of traditional techniques. This study focuses on the mathematical modeling of inorganic whole-blood DNA extraction, a fundamental traditional method, with the goal of understanding and optimizing parameters to improve its economic feasibility [7].

Traditional DNA extraction methodologies frequently employ organic solvents, such as phenol-chloroform-isoamyl alcohol (PCI), which are well-regarded for their effectiveness in producing high DNA yields, albeit at a high cost. In contrast, inorganic methods utilize more cost-effective materials like sodium chloride (NaCl). However, the optimization of inorganic-based extraction techniques remains an active area of research. Mathematical modeling serves as an essential tool in elucidating the complex interactions between inorganic materials and biological components, providing a systematic framework for predicting and manipulating outcomes. This study aims to develop an integrated model that synthesizes principles from chemistry, biology, statistics, and mathematics to optimize key parameters. Such an approach has the potential to revolutionize DNA extraction methods, particularly benefiting economically constrained applications in fields such as medicine, forensics, and genetic research [3].

Inorganic DNA extraction methodologies involve a diverse array of reagents and protocol parameters, including incubation temperature, duration, and proteinase K enzyme concentration, which are critical for enhancing DNA yield from whole blood samples. In this study, statistical approaches such as Response Surface Methodology (RSM) are employed to rigorously optimize these parameters. The primary objective is to determine the conditions that maximize DNA yield using a Box-Behnken response surface design. This investigation significantly contributes to defining the optimal conditions for the minimal necessary incubation temperature, duration, and proteinase K enzyme quantity, thereby ensuring the highest possible DNA yield [10].

## 2 Materials and Methods

### 2.1 Chemicals and reagent

DNA extraction is a cornerstone of molecular biology, essential for genetic research, forensic analysis, and medical diagnostics. The process employs various chemicals and reagents in a series of steps to isolate high-quality DNA. Initially, a lysis buffer, typically containing detergents like SDS (sodium dodecyl sulfate) or Triton X-100, disrupts the cell membrane to release DNA. Proteinase K is then used to degrade proteins, including histones, freeing the DNA from its associated proteins. Stabilization and precipitation of DNA are achieved using salts such as sodium chloride (NaCl) or potassium chloride (KCl). For further purification, organic solvents like phenol-chloroform or chloroform are employed to separate DNA from other cellular components. Finally, DNA is precipitated using ethanol or isopropanol and collected via centrifugation. These reagents work together to yield high-purity DNA suitable for downstream genetic analyses and biotechnological applications. In our study, whole blood was used as the starting material, with 6M NaCl effectively facilitating the precipitation of protein debris.

### 2.2 DNA extraction, quantification

Traditionally, DNA extraction from whole blood has relied predominantly on organic solvents and detergents, with limited exploration of purely inorganic methodologies. Inorganic approaches typically center on DNA precipitation utilizing salts or other inorganic agents after the initial cell lysis [2]. Presented below is a streamlined protocol employing inorganic precipitation for DNA extraction from whole blood. after the cell lysis with TE buffer, incubation with buffer A1, SDS, and proteinase K, A volume of 50 *µ*L 6M NaCl was vigorously agitated with the sample, followed by 15 minutes of incubation on ice. Subsequently, centrifugation at 4000 rpm for 15 minutes facilitated the sedimentation of salts and proteins. The resulting supernatant was carefully transferred to a freshly labeled 1.5-mL microcentrifuge tube. Addition of an equal volume of chilled isopropanol-induced DNA precipitation, visualized upon gentle inversion of the tubes. Notably, in our optimization endeavors, we have employed Response Surface Methodology with a Box-Behnken design to enhance DNA yield with the minimum number of experiments using Design Expert v8.06 software.

The process variables and their respective ranges for a given experiment are outlined in **Table 1**. These variables include Level, incubation temperature (°C), incubation time (hours), and proteinase K enzyme volume (µL). Each variable is categorized into three levels: –1, 0, and 1. For Temperature, the range spans from 54°C to 58°C. Time varies from 8 to 12 hours, while proteinase K enzyme volume ranges from 8 to 12 µL. This table serves as a reference for understanding the experimental conditions and their corresponding parameter ranges, crucial for ensuring consistency and repeatability in the experiment’s outcomes.

**Table 1:** Process Variables and Their Range.

| Values | Level | -1 | 0 | 1 |
| --- | --- | --- | --- | --- |
| A | Temperature($^{\circ}C$) | 54 | 56 | 58 |
| B | Time(hour) | 8 | 10 | 12 |
| C | Proteinasek($\mu L$) | 8 | 10 | 12 |

The experimental runs conducted for this purpose are outlined in **Table 2**. Through systematic exploration of these combinations, we conducted 17 experiments aimed at DNA yield from whole blood samples followed by DNA quantification.

**Table 2:** Results of Box Behnken Design.

| S.No. | A | B | C | Yield |
| --- | --- | --- | --- | --- |
| 1 | 56.00 | 10.00 | 10.00 | 350 |
| 2 | 56.00 | 8.00 | 8.00 | 90 |
| 3 | 56.00 | 10.00 | 10.00 | 350 |
| 4 | 54.00 | 10.00 | 8.00 | 110 |
| 5 | 56.00 | 10.00 | 10.00 | 350 |
| 6 | 58.00 | 10.00 | 12.00 | 310 |
| 7 | 56.00 | 8.00 | 12.00 | 270 |
| 8 | 56.00 | 10.00 | 10.00 | 350 |
| 9 | 56.00 | 10.00 | 10.00 | 350 |
| 10 | 54.00 | 10.00 | 12.0 | 100 |
| 11 | 58.00 | 10.00 | 8.00 | 180 |
| 12 | 58.00 | 12.00 | 10.00 | 300 |
| 13 | 56.00 | 12.00 | 8.00 | 150 |
| 14 | 54.00 | 12.00 | 10.00 | 220 |
| 15 | 54.00 | 8.00 | 10.00 | 200 |
| 16 | 56.00 | 12.00 | 12.00 | 400 |
| 17 | 58.00 | 8.00 | 10.00 | 180 |

## 3 Mathematical Modeling

### 3.1 Response surface methodology (RSM)

RSM is a mathematical technique utilized for modeling and analyzing diverse processes. In RSM, experiments examine the correlation between input variables (factors) and the output response. The objective is to enhance this response by adjusting the input variables within defined limitations. RSM designs typically entail conducting experiments at varied levels of the input variables, often guided by designs such as BBD or Central Composite designs (CCD). Particularly (BBD) facilitates efficient exploration of the factor space while minimizing the required number of experimental runs. Subsequently, statistical models, such as linear or quadratic regression models, are applied to the collected experimental data to characterize the relationship between the input variables and the response. These models serve to forecast optimal conditions for achieving the desired response of DNA yield. In this technique, the main objective is to optimize the response surface that is influenced by various process parameters. RSM also quantifies the relationship between the controllable input parameters and the obtained response surfaces [5].

### 3.2 Mathematical form of RSM design

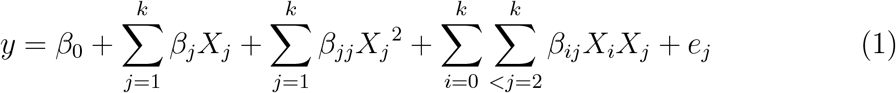

Let i, j=1,2,3 put in the equation.

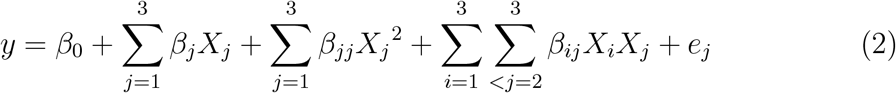

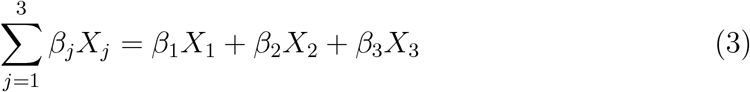

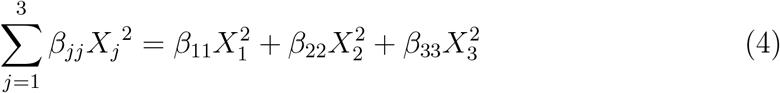

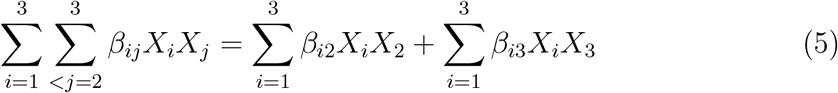

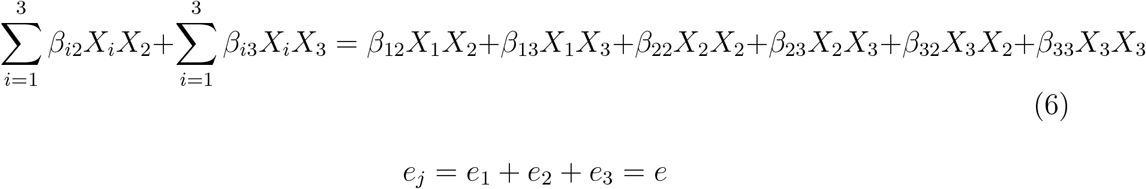

After putting all values in equation (2) and simplifying we get

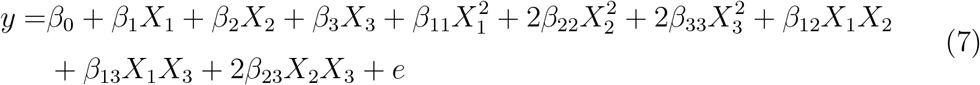

Equation (7) is possible when

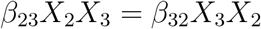

Now put *X*_1_ = *A, X*_2_ = *B*, and *X*_3_ = *C* in equation (7).

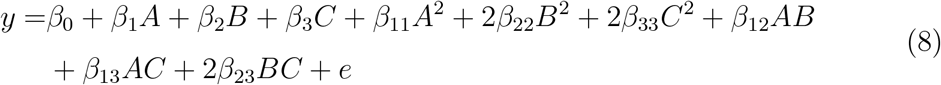

### 3.3 Box-bheken design (BBD)

BBD, commonly used in RSM, optimizes processes and analyzes the effects of multiple variables on a response. With three levels of each factor (−1, 0, +1), it explores linear and quadratic effects efficiently. This approach, requiring fewer runs than a full factorial design, reduces time and cost. Coupled with RSM, it fits models to experimental data, revealing complex relationships. BBD and RSM combined identify optimal process conditions, enhancing efficiency and quality in diverse fields.

### 3.4 Analysis of variance (ANOVA)

ANOVA compares means of three or more groups to detect significant differences by assessing within-group and between-group variability. It utilizes the F-statistic to evaluate this ratio, with a small p-value (*<* 0.05) and a large F-statistic indicating significant differences. Different types, such as one-way ANOVA, factorial ANOVA, and repeated measures ANOVA, serve to effectively evaluate group differences. The effectiveness of developed mathematical models was evaluated using Pareto ANOVA, with effective models utilized to create response surface contour graphs for examining variable interactions and response patterns.

## 4 Results

The equation represents the response function used to predict the extraction yield of DNA from whole blood. It incorporates various independent variables, namely incubation temperature (A), incubation time (B), and Proteinase K Enzyme Concentration (C), each with corresponding coefficients determined through experimental data analysis. The equation is structured as follows:

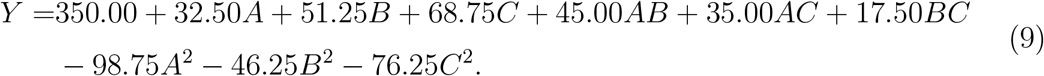

In this equation, (Y) represents the extraction yield of DNA in a microlitter. The coefficients accompanying each independent variable and their respective interactions reflect the impact of these factors on the extraction process. The quadratic terms (such as *A*^2^, *B*^2^, and *C*^2^) suggest potential nonlinear relationships between the variables and the extraction yield. This equation serves as a predictive tool for estimating DNA extraction yield based on the specified experimental conditions.

### 4.1 Effect of protocol variables on the extraction of DNA

Numerous protocol variables, such as incubation temperature, duration of incubation, and concentration of Proteinase K enzyme, play pivotal roles in DNA extraction from whole blood. These variables exhibit intricate interplay, significantly impacting the efficacy of DNA extraction. The provided equation encapsulates this intricate relationship through the allocation of coefficients to each variable and their interactions. Incubation temperature, duration, and enzyme concentration exert a direct influence on the yield of extracted DNA. Moreover, the inclusion of quadratic terms in the equation intimates that the correlation between these variables and DNA yield may not adhere strictly to linearity, suggesting the existence of optimal conditions for extraction. Appreciating the nuanced influence of these protocol variables on DNA extraction is indispensable for refining extraction protocols and achieving maximal DNA yields from whole blood specimens [8].

### 4.2 Effect of incubation temperature

The DNA extraction process from blood samples, whether stored at room temperature or subjected to 37°C incubation for 24 hours, yielded DNA quantities and quality akin to those obtained from samples cryopreserved at –70°C. Notably, samples undergoing more than four freeze-thaw cycles displayed discernible signs of partial DNA degradation [6].

### 4.3 Effect of DNA incubation time

Whole blood is a prominent DNA source, particularly in genotype diagnostic applications. Given the prevalent practice of receiving blood samples through mail or storing them before DNA extraction, comprehending the influence of storage duration (number of days elapsed from blood collection to DNA extraction) and temperature on DNA yield and quality is paramount. In our investigation, we meticulously evaluated DNA yield and quality derived from bovine blood samples subjected to diverse storage conditions, encompassing varying durations (3, 7, 14, or 28 days) and temperatures (−20°C, 4°C, 23°C, or 37°C), juxtaposed with the benchmark of DNA extraction performed within 4 hours of collection. Notably, the most robust mean DNA yields, relative to the control specimens, were attained from blood preserved at 4°C. Conversely, blood stored at 37°C for ≥ 3 days or 23°C for ≥7 days exhibited diminished DNA yields (P *<* .05) compared to counterparts stored at 4°C or –20°C for up to 28 days. The recovery of DNA was unsuccessful for three samples stored at 23°C and five samples stored at 37°C. Nevertheless, notwithstanding the diverse storage conditions, the genomic DNA extracted manifested high molecular weight and demonstrated suitability for restriction enzyme digestion and PCR amplification assays [4].

### 4.4 Effect of proteinase k enzyme concentration

Red blood cell (RBC) lysis, in conjunction with a two-step washing process, did not yield DNA of high purity, as evidenced by NanoDrop measurements. However, enhancements in DNA purity were achieved through the implementation of various modifications. These modifications encompassed elevating the concentration of proteinase K, prolonging the incubation period at 50°C, and introducing three washing steps during RBC lysis. These adjustments had a positive impact on the overall purity of the extracted DNA from the samples [9].

### 4.5 Multi response optimization and validation

Multi-response optimization and validation involve the process of simultaneously optimizing multiple responses or outcomes while ensuring the validity and reliability of the results. This approach allows for efficient exploration of various factors and their interactions to achieve desired objectives. By considering multiple responses, such as quality, cost, and efficiency, researchers can identify optimal conditions that balance these factors. Validation ensures that the optimized conditions are robust and reproducible across different experimental conditions, ensuring the reliability of the optimization process. Overall, multi-response optimization and validation methodologies enable comprehensive and effective decision-making in various fields, from manufacturing to experimental design.

## 5 Results of RSM Design using BBD

**Table 3:** This table provides a thorough overview of the sequential model sums of squares derived from the experimental design, encompassing the responses under investigation. It is structured into two distinct sections: the initial segment delineates the sequential model sums of squares of the total yield, comprising the sum of squares, degrees of freedom, mean square, F-value, and associated probability value for each model term, accompanied by pertinent remarks. Subsequently, the second segment furnishes model summary statistics, comprising the standard deviation, R-squared, adjusted R-squared, predicted R-squared, and PRESS (predicted residual sum of squares), along with corresponding remarks for each model term. These statistical metrics play a pivotal role in assessing the significance and efficacy of the linear, 2FI (two-factor interaction), quadratic, and cubic terms in elucidating the variability observed in the response variable. Such insights facilitate model refinement, aiding in its selection and interpretation within the experimental context.

**Table 3:** Sequential Model Sum of Squares for Responses.

| Source | Sum of Squares | df | Mean Square | F Value | Prob > F | Remarks |
| --- | --- | --- | --- | --- | --- | --- |
| Sequential model sum of squares for total yield |  |  |  |  |  |  |
| Mean | 1.028E+006 | 1.00 | 1.028E+006 |  |  |  |
| Linear | 67275.00 | 3.00 | 22425.00 | 2.43 | 0.1118 | Suggested |
| 2F1 | 14225.00 | 3.00 | 4741.67 | 0.45 | 0.7238 |  |
| Quadratic | 82336.76 | 3.00 | 27445.59 | 8.22 | 0.0108 | Suggested |
| Cubic | 23375.00 | 3.00 | 7791.67 | 6.366E+007 | <0.0001 | Aliased |
| Residual | 0.000 | 4.00 | 0.000 |  |  |  |
| Total | 1.21E+006 | 17.00 | 71470.59 |  |  |  |
| Model Summary Statistic |  |  |  |  |  |  |
| Source | Std.Dev | R-Squared | Adjusted R-Squared | Predicted R-Squared | PRESS | Remarks |
| Linear | 96.05 | 0.3594 | 0.2115 | -0.0618 | 1.988E+005 |  |
| 2F1 | 102.82 | 0.4353 | 0.0965 | -0.7405 | 3.258E+005 |  |
| Quadratic | 57.79 | 0.8751 | 0.7146 | -0.9977 | 3.740E+005 | Suggested |
| Cubic | 0.000 | 1.0000 | 1.00000 |  | + | Aliased |

**Table 4:** The ANOVA table presented herein encapsulates the analysis of variance about a Response Surface Quadratic Model, employed for discerning the interrelation between multiple independent variables denoted as A, B, and C, and a designated response variable. This tabular representation delineates various sources of variation encompassing the model itself, individual factors (A, B, C), interactions among factors (AB, AC, BC), and quadratic terms (*A*^2^, *B*^2^, *C*^2^). The “Sum of Squares” column quantifies the extent of variability in the response variable attributed to each factor or combination thereof. Degrees of freedom (Df) denote the number of independent pieces of information utilized for parameter estimation. The “Mean Square” represents the ratio of the sum of squares to its corresponding degrees of freedom. The “F Value” serves as a metric gauging the ratio of variability among group means to variability within the groups, facilitating the assessment of each factor’s significance. The “p-value” delineates the probability of observing a given F value or larger under the null hypothesis, with a lower p-value (typically *<*0.05) indicative of statistical significance for the factor or interaction under consideration. Supplementary metrics such as the Coefficient of Variation (CV), Prediction Sum of Squares (PRESS), and R-squared values furnish additional insights into the model’s efficacy and predictive prowess.

**Table 4:** ANOVA Result for Response.

| Source | Sum of Squares | Df | Mean Square | F Value | P Value<br>Prob > F |
| --- | --- | --- | --- | --- | --- |
| Model | 1.638E+005 | 9 | 18204.08 | 5.45 | 0.0180 |
| A-Temperature | 8450.00 | 1 | 8450.00 | 2.53 | 0.1557 |
| B-Time | 21012.50 | 1 | 21012.50 | 6.29 | 0.0405 |
| C-Proteinase k | 37812.50 | 1 | 37812.50 | 11.32 | 0.0120 |
| AB | 8100.00 | 1 | 8100.00 | 2.43 | 0.1633 |
| AC | 4900.00 | 1 | 4900.00 | 1.47 | 0.2651 |
| BC | 1225.00 | 1 | 1225.00 | 0.37 | 0.5638 |
| $A^2$ | 41059.21 | 1 | 41059.21 | 12.30 | 0.0099 |
| $B^2$ | 9006.58 | 1 | 9006.58 | 2.70 | 0.1445 |
| $C^2$ | 24480.26 | 1 | 24480.26 | 7.33 | 0.0303 |
| CV | 23.50 |  |  |  |  |
| AP | 5.895 |  |  |  |  |
| PRESS | 3.740+005 |  |  |  |  |
| Mean | 245.88 |  |  |  |  |
| $R^2$ | 0.8751 | | | | |
| Adj- $R^2$ | 0.7146 | | | | |

**Figure 1:**
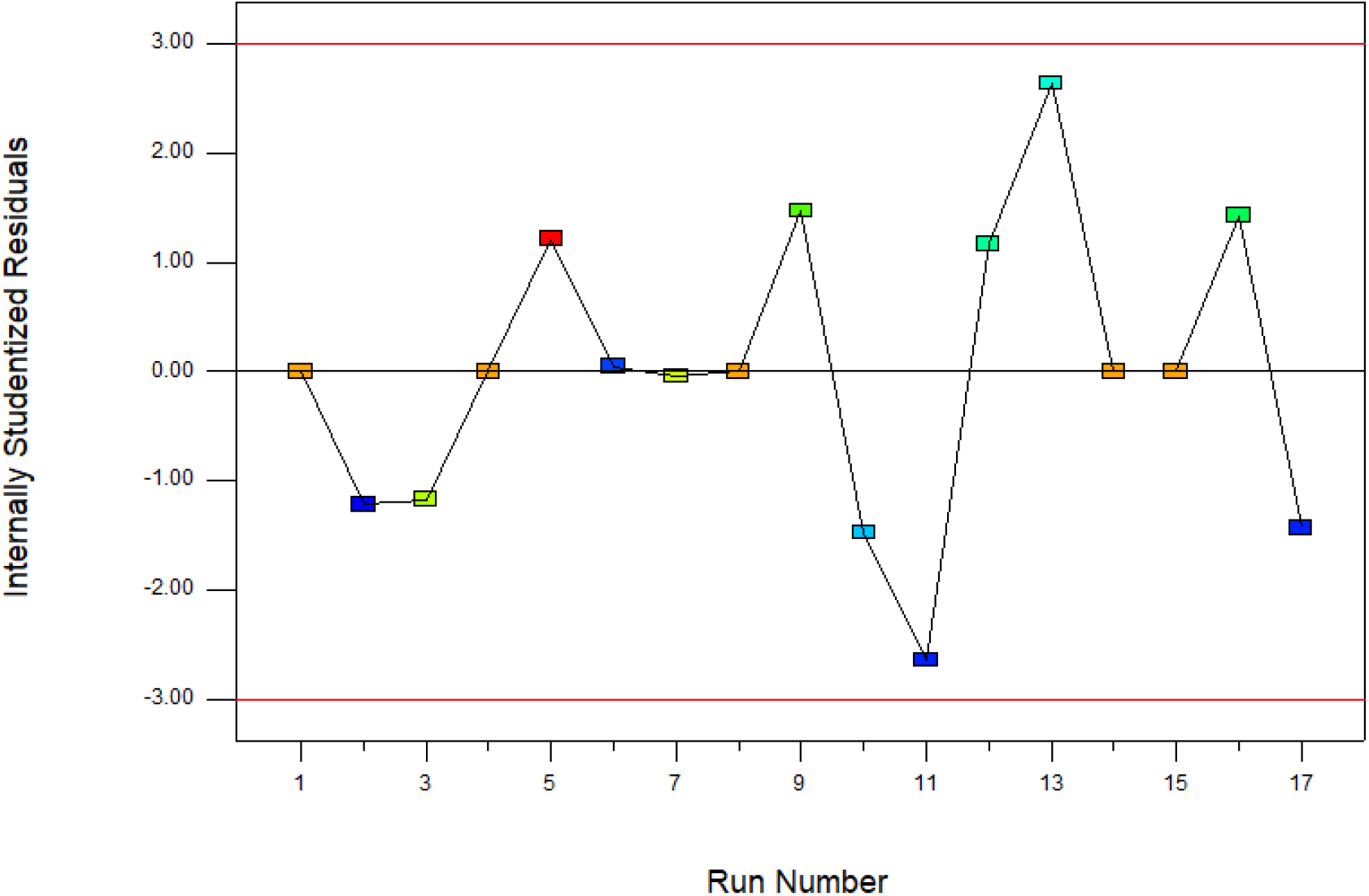
The statistical analysis revealed a highly significant model F value along with a meager probability value (0.0001), indicating the robustness of the model and its capability to effectively describe the response. A normal distribution function was employed to fit the studentized residuals. Subsequently, a comparison was made between the predicted studentized residuals derived from the best-fit normal distribution and the experimentally obtained studentized residuals.

**Figure 2:**
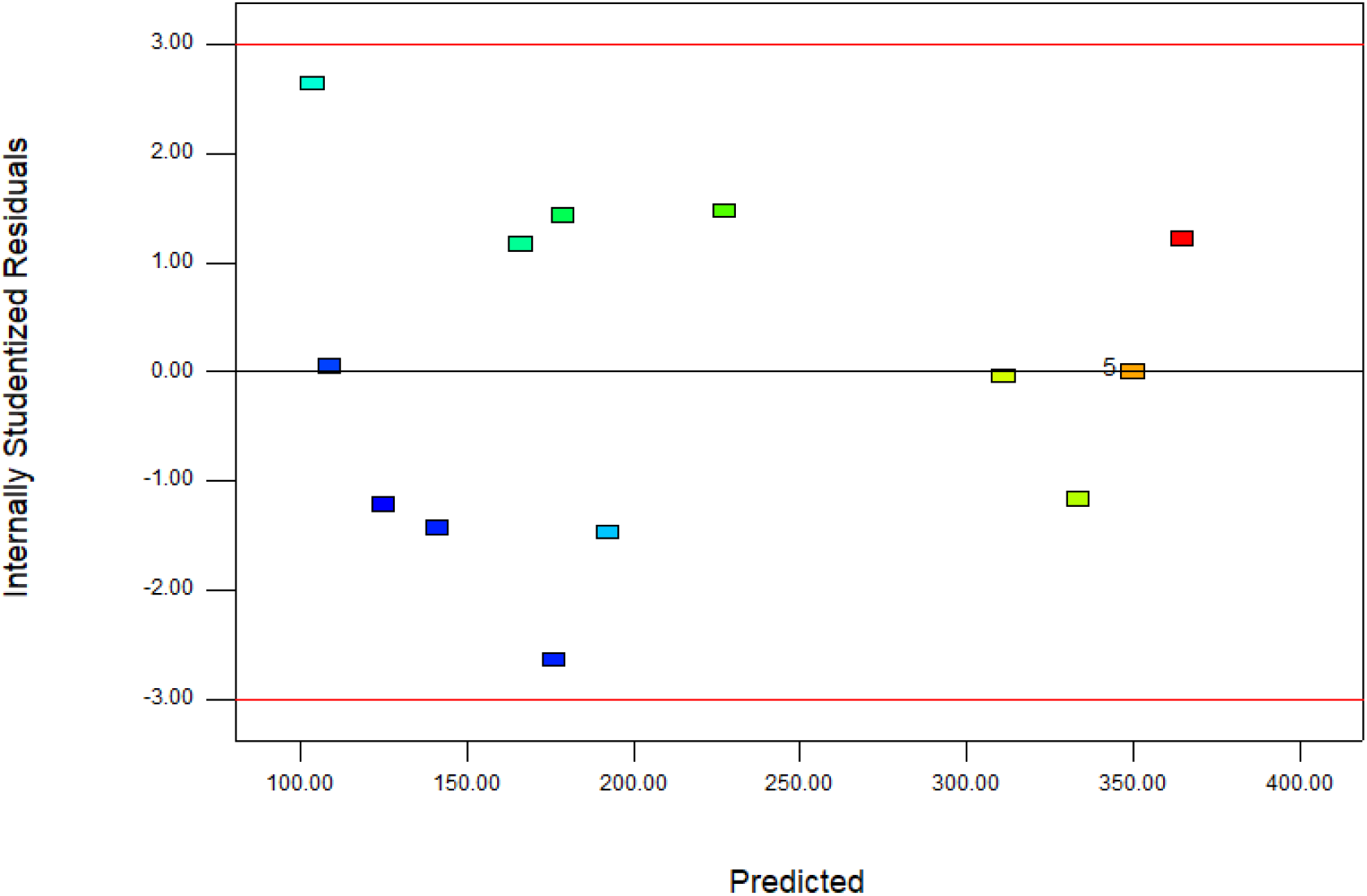
Moreover, diagnostic plots like the predicted versus actual plot aid in assessing the model’s appropriateness and elucidating the connection between predicted and experimental values. The proximity of data points to the straight line in this plot suggests a satisfactory agreement between actual data and the model-derived data.

**Figure 3:**
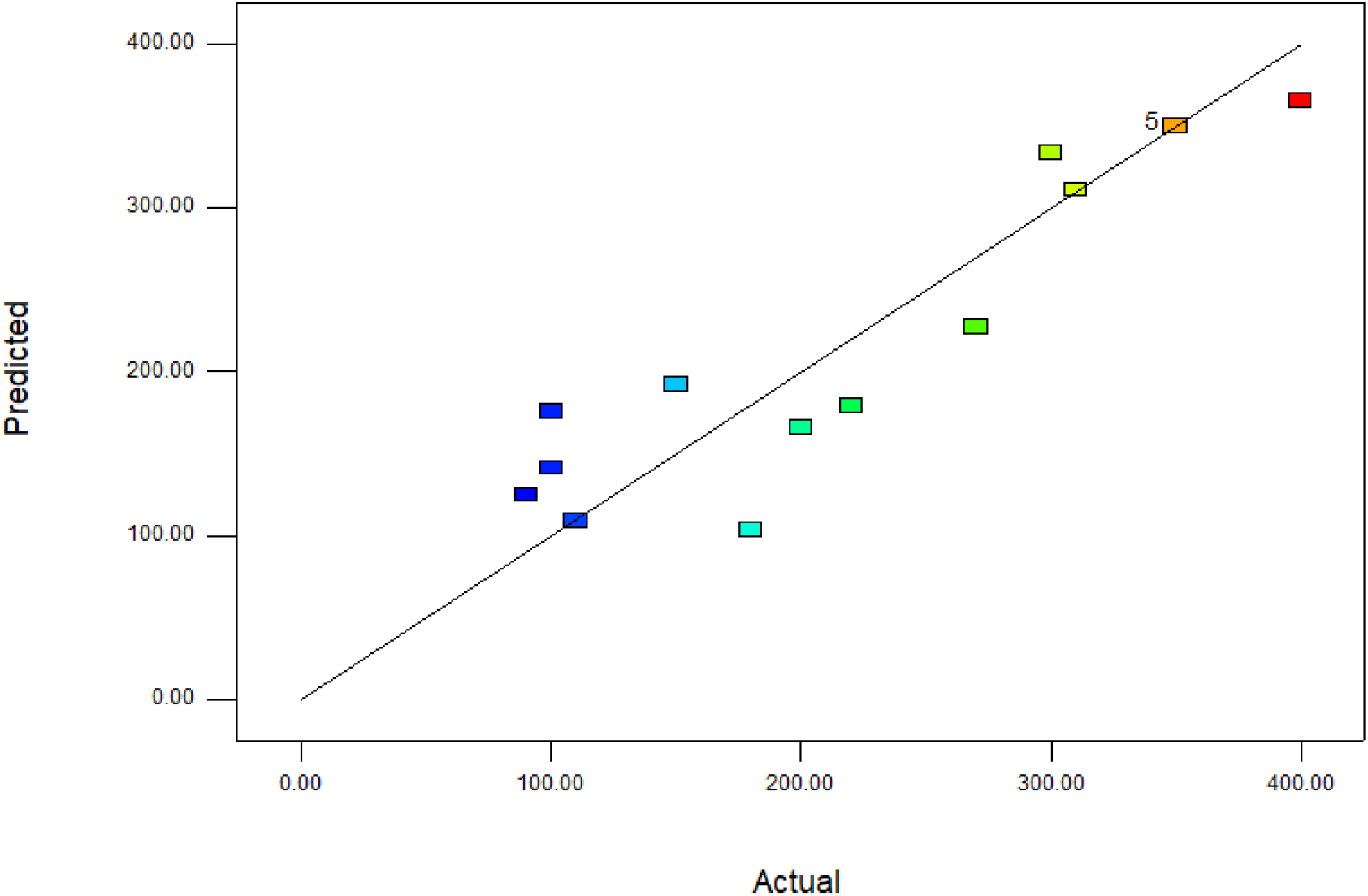
Shows a plot of the studentized residuals against their predicted values is depicted. A random scatter pattern in this plot is anticipated, suggesting that the fluctuations in the original observations are independent of the response values. Such a pattern further reinforces the appropriateness of the proposed model in describing the underlying process.

**Figure 4:**
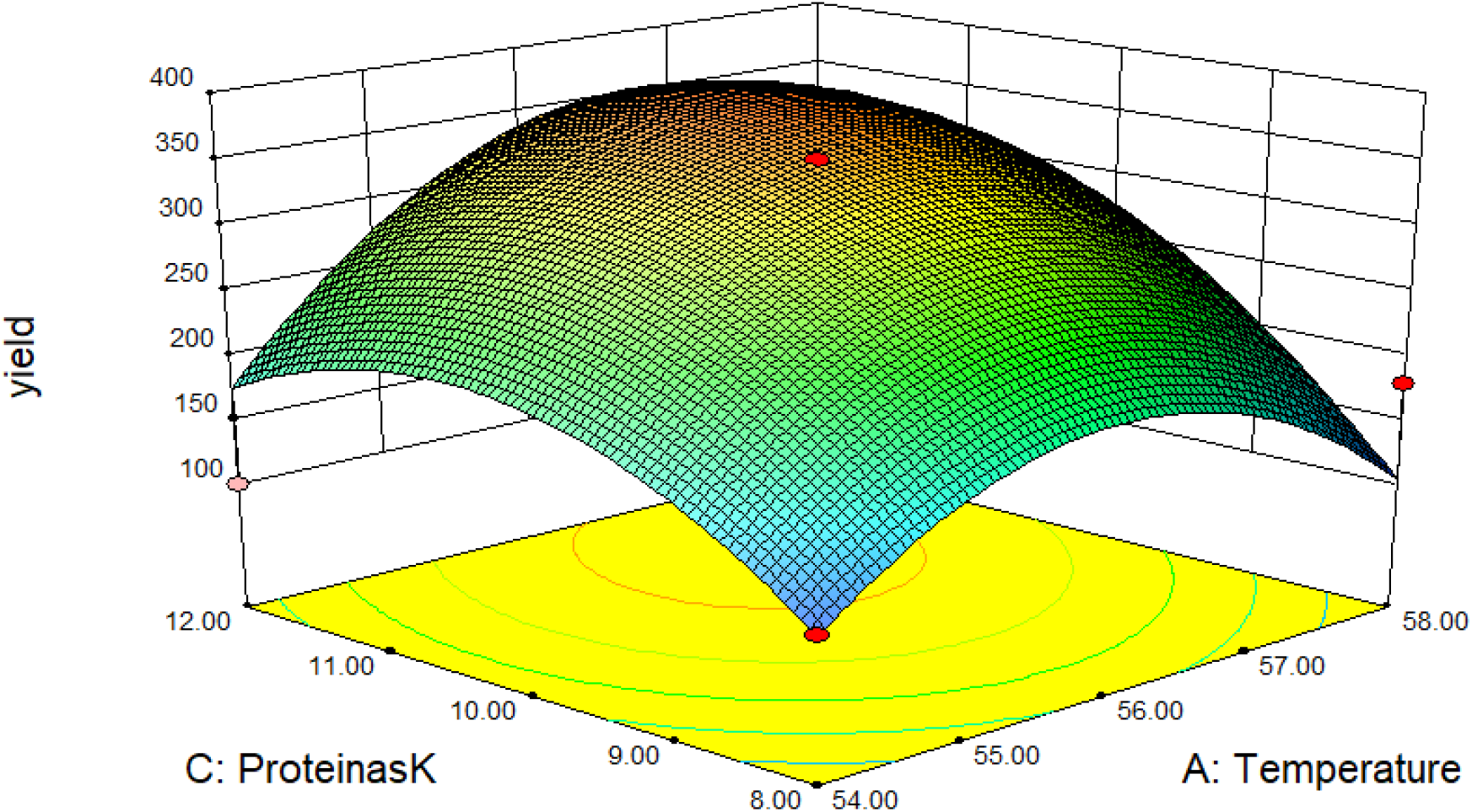
The plot depicts a three-dimensional surface against a grid background, demonstrating how variations in protein concentration and temperature influence yield. This 3D surface plot effectively shows the impact of protein concentration and temperature on yield.

## 6 Conclusion

In the current investigation, enzyme-assisted DNA extraction was conducted to optimize and assess the impact of different parameters, including incubation temperature, incubation time, and Proteinase K enzyme concentration, on the DNA yield. Response Surface Methodology (RSM) combined with Box-Behnken Design (BBD) was utilized to analyze and optimize the process variables influencing the extraction DNA yield. The outcomes revealed significant effects of all process variables on the DNA yield, leading to the development of a quadratic model for response prediction. The optimal conditions resulting in the highest DNA yield (300*µ*L) were determined as follows: incubation temperature of 56 °C, incubation time of 10 hours, and a proteinase K enzyme concentration of 10*µ*L.

## References

[1] André I Khuri and Siuli Mukhopadhyay, 2010: Response surface methodology. Wiley Interdisciplinary Reviews: Computational Statistics, 2(2):128–149.

[2] Ashok Kumar Dogra and Archana Prakash, 2023: An Effective and Rapid Method of DNA Extraction Protocol from Samples of Human Blood. 12, 10.5530/ajbls.2023.12.25..

[3] Crooke, P. S., and F. F. Parl, 2010: A Mathematical Model for DNA Damage and Repair. 2010, 10.4061/2010/352603.

[4] Cushwa, W., and J. F. Medrano, 1993: Effects of blood storage time and temperature on DNA yield and quality. 14.

[5] Gilmour, S., 2006: Response Surface Designs for Experiments in Bioprocessing. 62, 10.1111/J.1541-0420.2005.00444.X.

[6] Lahiri, D. K., and B. Schnabel, 1993: DNA isolation by a rapid method from human blood samples: Effects of MgCl2, EDTA, storage time, and temperature on DNA yield and quality. 31, 10.1007/BF02401826.

[7] Lindahl, T., 1993: Instability and decay of the primary structure of DNA. 362, 10.1038/362709A0.

[8] Martínez de la Puente, J., S. Ruiz, R. C. Soriguer, and J. Figuerola, 2013: Effect of blood meal digestion and DNA extraction protocol on the success of blood meal source determination in the malaria vector Anopheles atroparvus. 12, 10.1186/1475-2875-12-109.

[9] Shahriar, M., R. Haque, S. Kabir, I. Dewan, and M. A. Bhuyian, 2011: Effect of Proteinase-K on Genomic DNA Extraction from Gram-positive Strains. 4, 10.3329/SJPS.V4I1.8867.

[10] Silva, E., H. Rogez, and Y. Larondelle, 2007: Optimization of extraction of phenolics from inga edulis leaves using response surface methodology. 55, 381–387.

